# Declining relative humidity in the Thompson River Valley near Kamloops, British Columbia, Canada: 1990-2012

**DOI:** 10.1101/028894

**Authors:** Sierra Rayne, Kaya Forest

## Abstract

Potential time trends in relative humidity (RH) were investigated for the Kamloops climate station in a semi-arid region of south-central British Columbia, Canada, between 1990 and 2012. Mean monthly 6 am and 3 pm RH at Kamloops achieve annual minima during the March to September period with substantially higher early morning RH compared to the mid-afternoon period. Significant temporal declines in RH throughout the year are evident ranging from 1.5 to 5.7%/decade. No significantly increasing temporal trends in RH were found. The findings indicate that a continuation of declining trends in RH for the study area may increase the quantity of dust and other atmospheric particulate generation from both natural and anthropogenic sources, possibly resulting in additional threats to local and regional air quality, thereby necessitating inclusion in air quality management planning and modeling efforts.

## Introduction

Relative humidity (RH) is an important atmospheric determinant for the distribution and occurrence of clouds, human comfort, and water and air quality management, as well as being integral to coherent agricultural, forestry, and other land use decision making efforts (Sundqvist 1978; Peixoto and Oort 1996; Price and Wood 2002; Wright et al. 2010). Along with wind speed, RH is also a dominant factor in determining rates of evaporation. Both observational trends and modeling studies have demonstrated that while increasing air temperatures over large open water bodies (e.g., marine systems) may result in relatively constant RH (i.e., increasing evaporation rates keep pace with increased water-holding capacity of the atmosphere), in evaporation constrained terrestrial regions rising air temperatures may lead to decreasing RH trends (Wetzel et al. 1996; Betts 2000; Redelsperger et al. 2002; Derbyshire et al 2004; Luo and Rossow 2004; Seager et al. 2007; Pierrehumbert et al. 2007; Betts and Ridgway 1988; Sherwood 2010; Sherwood et al. 2010; Risi et al. 2012; Ruosteenoja and Raisanen 2012; Fischer and Knutti 2013). However, substantial regional heterogeneity in the existence, direction, and magnitude of RH temporal trends is evident (Solomon et al. 2007; Willett 2007), with reports of declining RH trends over the global and hemispheric oceans (Willett 2007; Dai 2006), either no significant trends (Dai 2006) or increasing trends (Willett 2007) over the Northern Hemisphere land areas, declining trends over the Southern Hemisphere land areas (Willett 2007), and no significant trends over the global land mass (Willett 2007; Dai 2006).

A concern surrounding potential climate change impacts on RH temporal trends involves possible increasing risks to local and regional air quality from dust generation, particularly those from anthropogenic activities such as surface mining in arid and semi-arid systems which may generate air particulate having negative health impacts on nearby ecosystems and human populations (see, e.g., Martin et al. 1975; Le Bouffant et al. 1982; Khan et al. 1983; Schins and Borm 1999; Branquinho et al. 1999; Reynolds et al. 2003; Fubini and Fenoglio 2007; Corriveau et al. 2011). For example, in the semi-arid Thompson River Valley of south-central British Columbia, Canada, near the city of Kamloops (Figure 1), there are significant concerns among the local community regarding proposed mining activities near populated areas. This semi-arid region already experiences periodic episodes of poor air quality due to local industrial emissions, natural dust releases during dry and windy weather, and thermal inversions. Reduced RH will not only increase the air quality risks from existing dust sources, but may also act to increase dust generation from any nearby mining developments. In order to better understand these air quality issues, in the current work we investigate potential time trends in RH, as well as other related climate variables, for the Kamloops climate station between 1990 and 2012.

**Figure 1.**
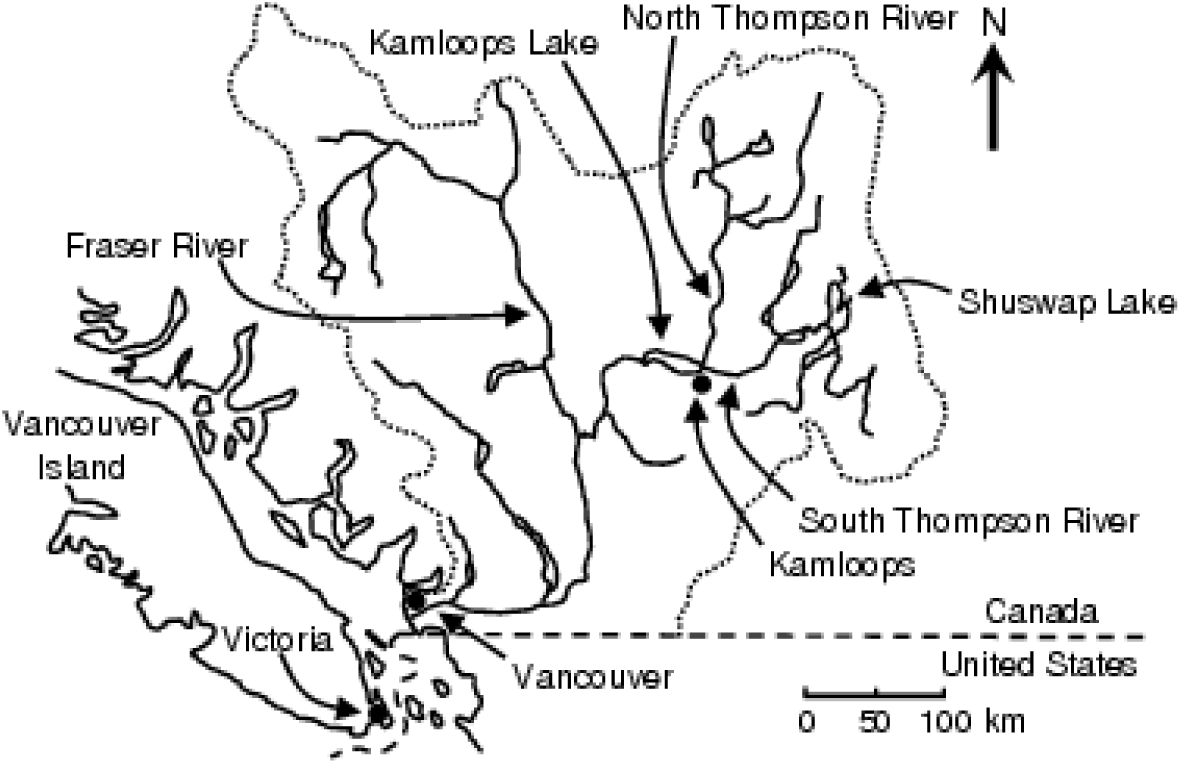
Location of the study area in southern British Columbia, Canada (adapted from www.ilec.or.jp/database/nam/nam-54.html.)

## Data and Analysis

Climate data for hourly relative humidity and monthly extreme maximum temperatures, extreme minimum temperatures, end of month total snow on ground, and direction and speed of maximum monthly wind gusts at the Kamloops A climate station (latitude: 50°42’08.000” N; longitude: 120°26’31.000” W; elevation: 345.30 m; climate ID: 1163780; WMO ID: 71887) in south-central British Columbia, Canada, were obtained from the online National Climate Data and Information Archive of Environment Canada (climate.weatheroffice.gc.ca). Monthly means of daily mean temperature, daily maximum temperature, and daily minimum temperature, as well as monthly total precipitation, rainfall, snowfall, and monthly mean of the hourly wind speeds at the Kamloops A climate station were taken from the second generation homogenized temperature, the second generation adjusted precipitation, and the homogenized surface wind speed datasets of the Adjusted and Homogenized Canadian Climate Data archive (Vincent et al. 2012; Mekis and Vincent 2011; Wan et al. 2009; Wang 2008).

Parametric linear regression and Spearman/Kendall non-parametric rank correlation analyses were conducted with KyPlot v.2.0.b.15. For continuous time series, trends were also examined using the non-parametric Mann-Kendall method with the Sen’s slope (Mann 1945; Kendall 1975; Sen 1968) within the R statistical software package (R Core Team 2012).

## Results and Discussion

Although a complete annual series of monthly RH data is available at Kamloops since 1953 (Figures 2 through 5), we restricted our temporal studies to the 1990 to 2012 period in order to avoid potential spurious trend analyses resulting from inclusion of the period during the 1970s through 1980s when the replacement of the psychrometer by the dewcel often created an artificial negative step in annual RH time series (van Wijngaarden and Vincent 2005; Vincent et al. 2007). Efforts can be undertaken to account for this possible discontinuity via interstation homogenization. However, given the heterogeneity in regional climates and land use changes over the past several decades within southcentral British Columbia, we viewed such correction measures as potentially unreliable and instead chose to focus on the post-1990 period free from any apparent instrument biases.

**Figure 2.**
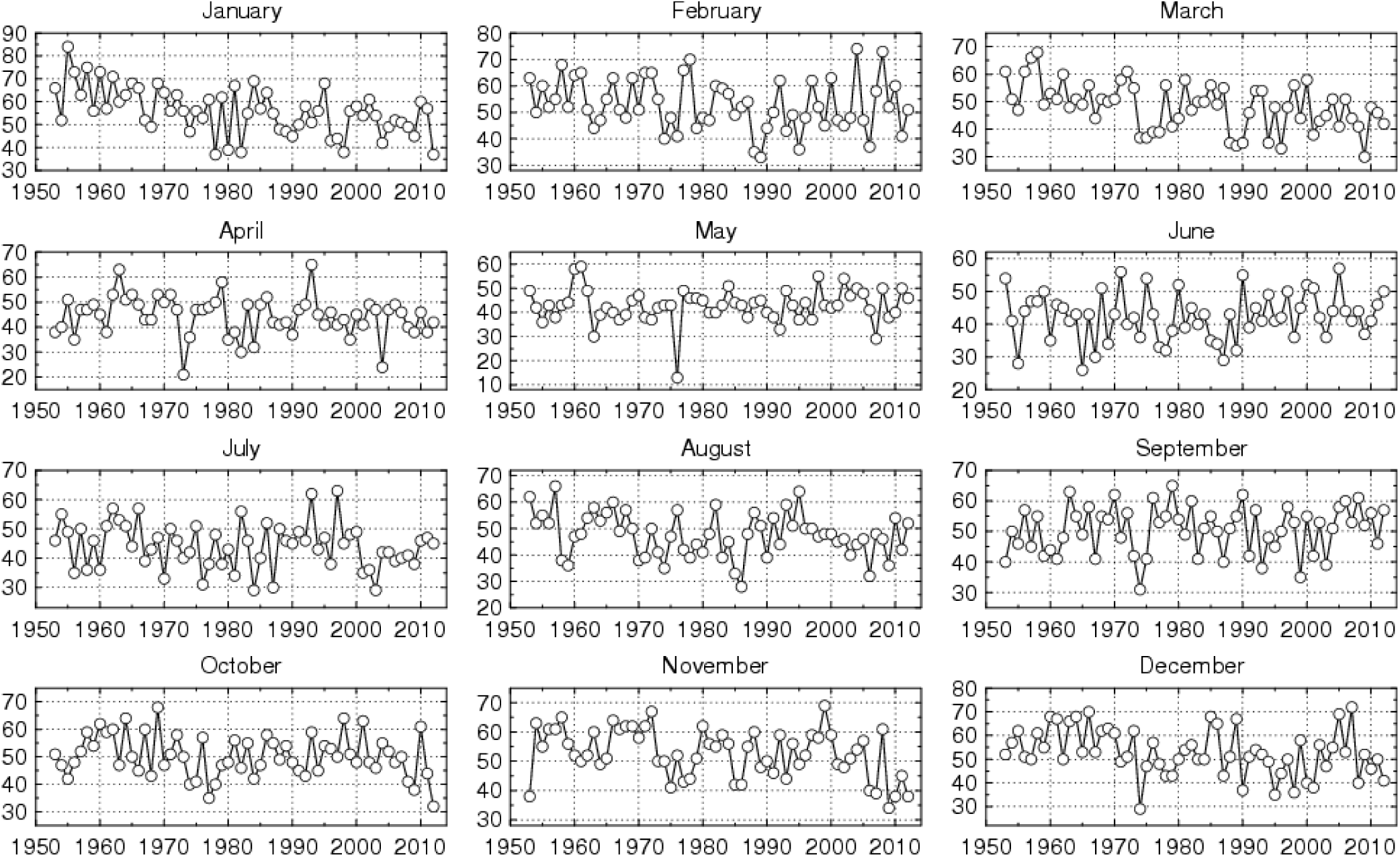
Monthly 6 am minimum relative humidity at the Kamloops A climate station between 1953 and 2012.

**Figure 3.**
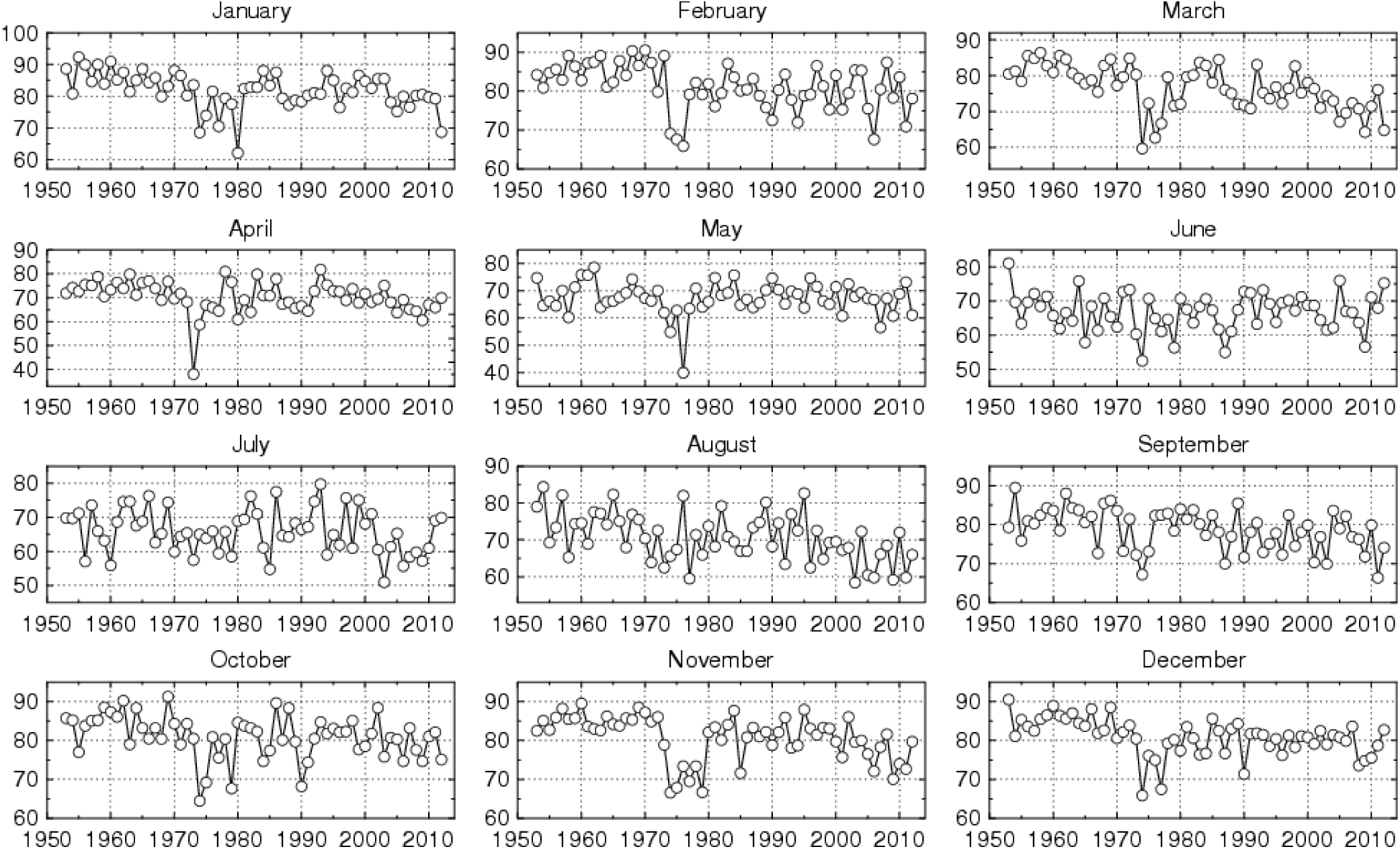
Monthly 6 am average relative humidity at the Kamloops A climate station between 1953 and 2012.

**Figure 4.**
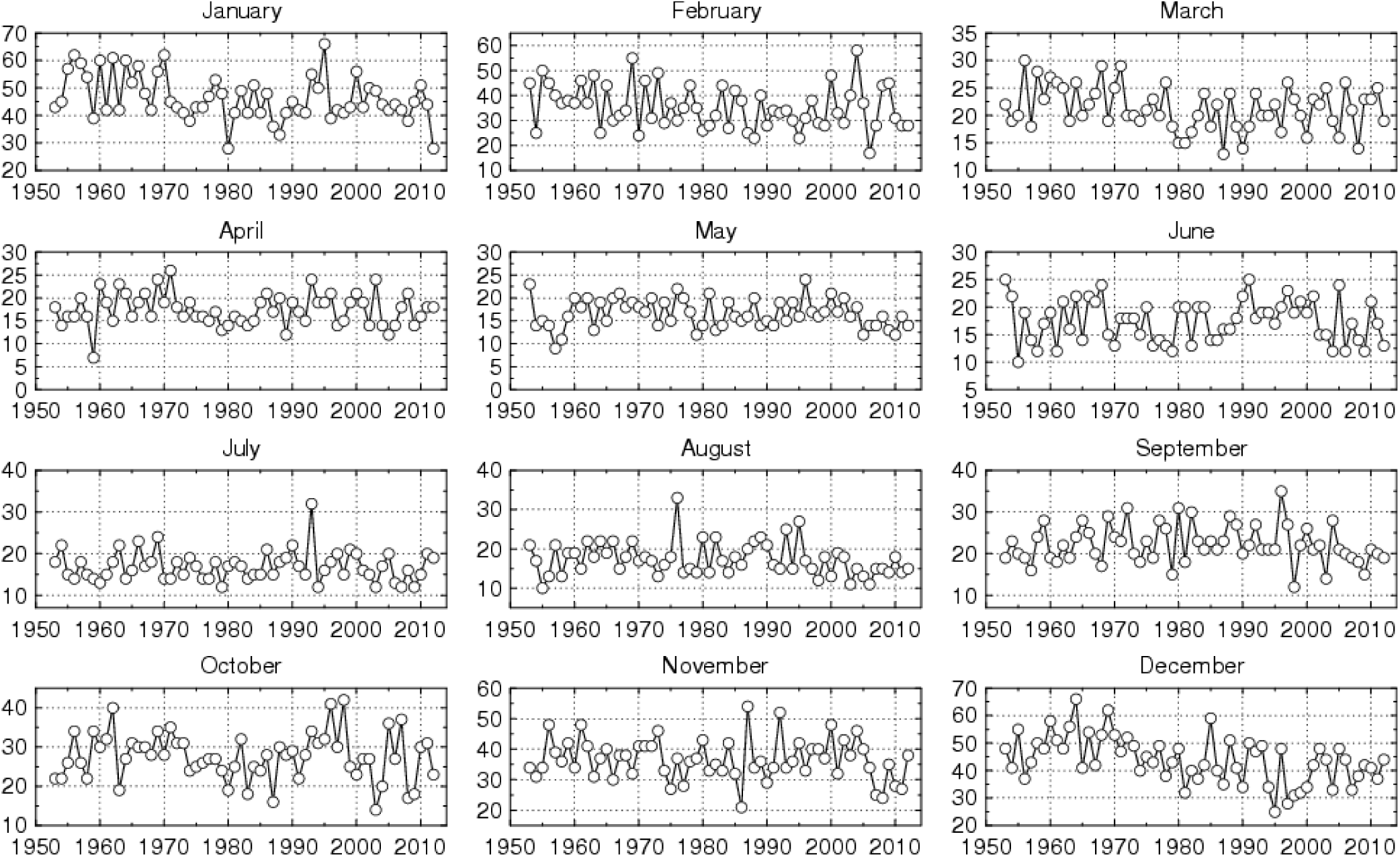
Monthly 3 pm minimum relative humidity at the Kamloops A climate station between 1953 and 2012.

**Figure 5.**
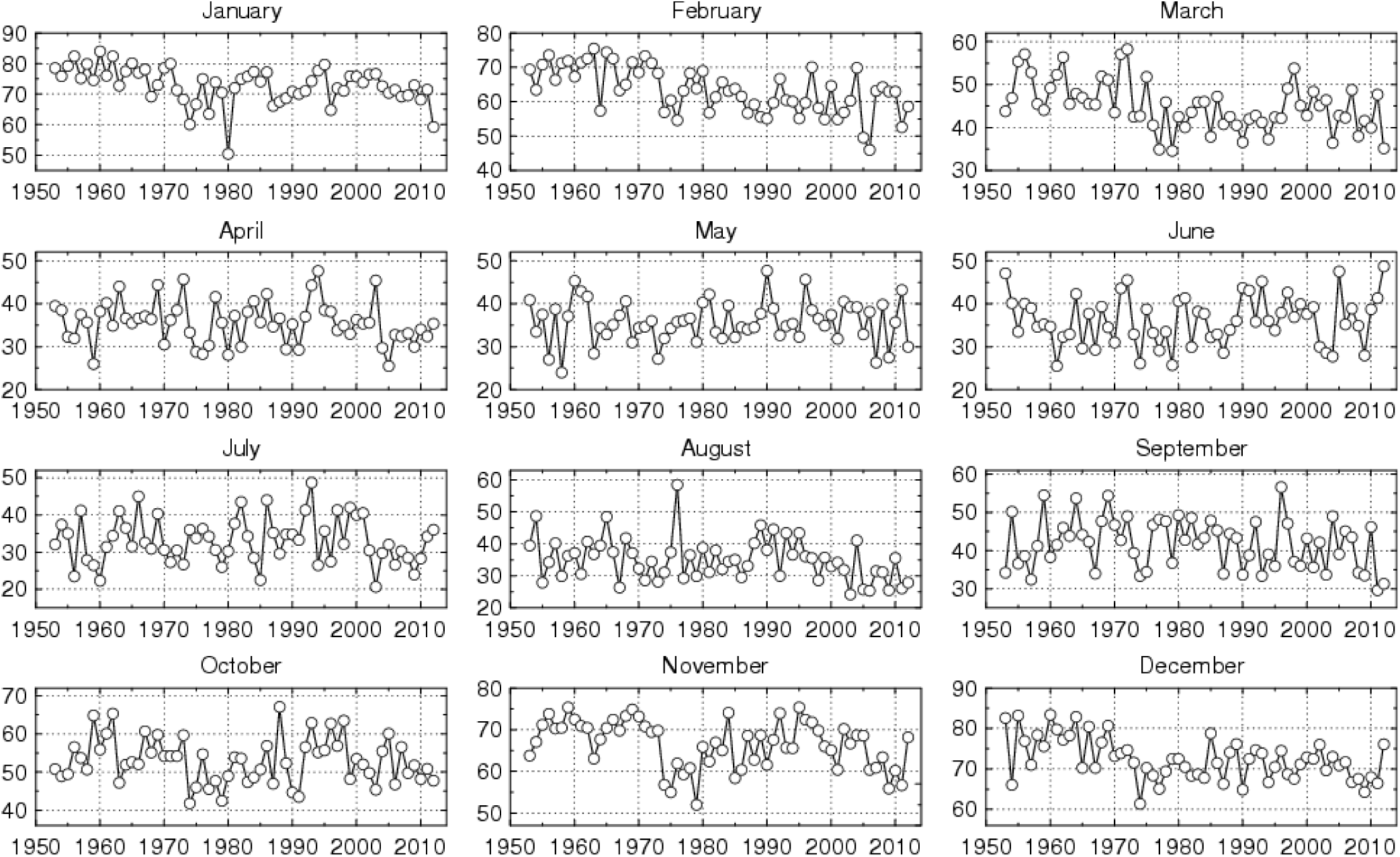
Monthly 3 pm average relative humidity at the Kamloops A climate station between 1953 and 2012.

Mean monthly 6 am (06:00:00h) and 3 pm (15:00:00h) RH at Kamloops between 1990 and 2012 achieve annual minima during the March to September period with substantially higher early morning RH compared to the mid-afternoon period (Table 1). Over this timeframe, we find significant (p≤0.1) temporal declines in RH throughout the year ranging from 1.5 to 5.7%/decade. No significantly increasing temporal trends in RH were found. Excellent agreement in trend significance and magnitude was observed between the parametric linear regression and non-parametric Mann-Kendall statistical methods. Our results are in agreement with those of Vincent et al. (2007) who reported significant declines in RH across western and southern Canada between 1953 and 2012.

**Table 1.** Mean values and parametric/non-parametric (Mann-Kendall) temporal trends in monthly relative humidity at the Kamloops A climate station between 1990 and 2012. Mean values are in units of %. Corresponding temporal trends are on a %/yr basis.

| Descriptor | Jan | Feb | Mar | Apr | May | Jun | Jul | Aug | Sep | Oct | Nov | Dec |
| --- | --- | --- | --- | --- | --- | --- | --- | --- | --- | --- | --- | --- |
| 6 am minimum relative humidity |  |  |  |  |  |  |  |  |  |  |  |  |
| Mean | 51.2 | 51.6 | 44.8 | 43.5 | 43.3 | 44.7 | 44.1 | 47.2 | 50.9 | 49.5 | 50.2 | 48.9 |
| Trend |  |  |  |  |  |  |  |  |  |  |  |  |
| Parametric | NS | NS | NS | NS | NS | NS | -0.40/+ | NS | NS | NS | -0.56/* | NS |
| Mann-Kendall | NS | NS | NS | NS | NS | NS | NS | NS | NS | NS | -0.55/+ | NS |
| 6 am average relative humidity |  |  |  |  |  |  |  |  |  |  |  |  |
| Mean | 80.7 | 79.1 | 73.4 | 69.4 | 67.6 | 67.9 | 65.0 | 67.6 | 76.1 | 79.7 | 79.5 | 79.3 |
| Trend |  |  |  |  |  |  |  |  |  |  |  |  |
| Parametric | -0.26/+ | NS | -0.35/* | -0.33/* | -0.25/* | NS | -0.40/+ | -0.42/* | NS | NS | -0.37/** | NS |
| Mann-Kendall | NS | NS | -0.36/* | -0.39/* | NS | NS | NS | -0.40/* | NS | NS | -0.39/* | NS |
| 3 pm minimum relative humidity |  |  |  |  |  |  |  |  |  |  |  |  |
| Mean | 45.2 | 33.7 | 20.7 | 17.6 | 16.3 | 18.1 | 17.2 | 16.2 | 21.4 | 28.0 | 36.3 | 39.4 |
| Trend |  |  |  |  |  |  |  |  |  |  |  |  |
| Parametric | NS | NS | NS | NS | -0.17/+ | -0.31/** | NS | -0.25/* | NS | NS | NS | NS |
| Mann-Kendall | NS | NS | NS | NS | -0.15/+ | -0.32/* | NS | -0.18/+ | -0.17/* | NS | NS | NS |
| 3 pm average relative humidity |  |  |  |  |  |  |  |  |  |  |  |  |
| Mean | 80.0 | 59.4 | 43.0 | 35.2 | 36.5 | 37.9 | 33.2 | 33.3 | 39.6 | 53.0 | 65.9 | 70.5 |
| Trend |  |  |  |  |  |  |  |  |  |  |  |  |
| Parametric | -0.23/+ | NS | NS | -0.29/+ | NS | NS | -0.41/+ | -0.57/** | NS | NS | -0.41/* | NS |
| Mann-Kendall | NS | NS | NS | -0.26/+ | NS | NS | -0.35/+ | -0.52/** | NS | NS | -0.44/* | NS |

NS= $p > 0.1$ ; += $0.05 < p \leq 0.1$ ; \*= $0.01 < p \leq 0.05$ ; \*\*= $0.001 < p \leq 0.01$ ; and \*\*\*= $p \leq 0.001$ .

Mean daily maximum temperatures have increased during mid-summer (July and August) and early autumn (October) at Kamloops between 1990 and 2011 at rates ranging from 0.65 to 1.3°C/decade (Table 2). No trends in mean daily temperatures or extreme maximum temperatures were observed. Isolated declines in mean daily minimum temperatures (April; -0.64°C/decade [parametric; the corresponding non-parametric trend was not significant]) and extreme minimum temperatures (April [- 1.1°C/decade (non-parametric; the corresponding parametric trend was not significant)] and July [-1.2/-1.3°C/decade]) are also evident. Total monthly precipitation and rainfall are in decline during June, July, and August (Table 3) by 11 to 16 mm/decade, indicating -- when coupled with increasing trends for daily maximum temperatures over this period -- a likely general increase in summertime aridity for the study area. No significant trends were found for monthly snowfall totals or end of month total snow on the ground. December has a significantly declining trend for total precipitation and rainfall, but no trend for snowfall.

**Table 2.**
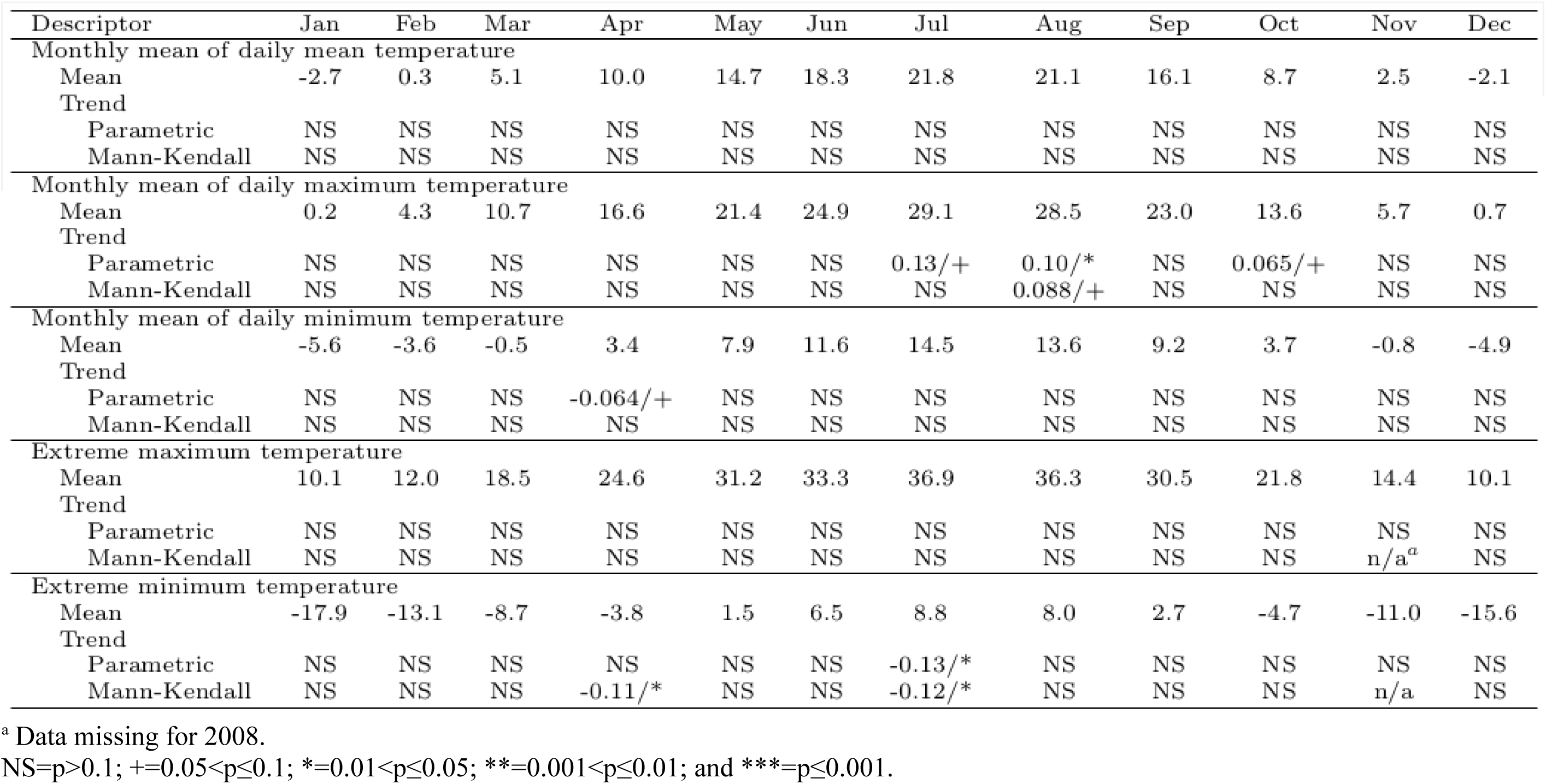
Mean values and parametric/non-parametric (Mann-Kendall) temporal trends in monthly daily mean, daily maximum, daily minimum, extreme daily maximum, and extreme daily minimum temperatures at the Kamloops A climate station between 1990 and 2011. Mean values are in units of °C. Corresponding temporal trends are on a °C/yr basis.

| Descriptor | Jan | Feb | Mar | Apr | May | Jun | Jul | Aug | Sep | Oct | Nov | Dec |
| --- | --- | --- | --- | --- | --- | --- | --- | --- | --- | --- | --- | --- |
| Monthly mean of daily mean temperature |  |  |  |  |  |  |  |  |  |  |  |  |
| Mean | -2.7 | 0.3 | 5.1 | 10.0 | 14.7 | 18.3 | 21.8 | 21.1 | 16.1 | 8.7 | 2.5 | -2.1 |
| Trend |  |  |  |  |  |  |  |  |  |  |  |  |
| Parametric | NS | NS | NS | NS | NS | NS | NS | NS | NS | NS | NS | NS |
| Mann-Kendall | NS | NS | NS | NS | NS | NS | NS | NS | NS | NS | NS | NS |
| Monthly mean of daily maximum temperature |  |  |  |  |  |  |  |  |  |  |  |  |
| Mean | 0.2 | 4.3 | 10.7 | 16.6 | 21.4 | 24.9 | 29.1 | 28.5 | 23.0 | 13.6 | 5.7 | 0.7 |
| Trend |  |  |  |  |  |  |  |  |  |  |  |  |
| Parametric | NS | NS | NS | NS | NS | NS | 0.13/+ | 0.10/* | NS | 0.065/+ | NS | NS |
| Mann-Kendall | NS | NS | NS | NS | NS | NS | NS | 0.088/+ | NS | NS | NS | NS |
| Monthly mean of daily minimum temperature |  |  |  |  |  |  |  |  |  |  |  |  |
| Mean | -5.6 | -3.6 | -0.5 | 3.4 | 7.9 | 11.6 | 14.5 | 13.6 | 9.2 | 3.7 | -0.8 | -4.9 |
| Trend |  |  |  |  |  |  |  |  |  |  |  |  |
| Parametric | NS | NS | NS | -0.064/+ | NS | NS | NS | NS | NS | NS | NS | NS |
| Mann-Kendall | NS | NS | NS | NS | NS | NS | NS | NS | NS | NS | NS | NS |
| Extreme maximum temperature |  |  |  |  |  |  |  |  |  |  |  |  |
| Mean | 10.1 | 12.0 | 18.5 | 24.6 | 31.2 | 33.3 | 36.9 | 36.3 | 30.5 | 21.8 | 14.4 | 10.1 |
| Trend |  |  |  |  |  |  |  |  |  |  |  |  |
| Parametric | NS | NS | NS | NS | NS | NS | NS | NS | NS | NS | NS | NS |
| Mann-Kendall | NS | NS | NS | NS | NS | NS | NS | NS | NS | NS | n/a<sup>a</sup> | NS |
| Extreme minimum temperature |  |  |  |  |  |  |  |  |  |  |  |  |
| Mean | -17.9 | -13.1 | -8.7 | -3.8 | 1.5 | 6.5 | 8.8 | 8.0 | 2.7 | -4.7 | -11.0 | -15.6 |
| Trend |  |  |  |  |  |  |  |  |  |  |  |  |
| Parametric | NS | NS | NS | NS | NS | NS | -0.13/* | NS | NS | NS | NS | NS |
| Mann-Kendall | NS | NS | NS | -0.11/* | NS | NS | -0.12/* | NS | NS | NS | n/a | NS |

<sup>a</sup> Data missing for 2008.
NS= $p > 0.1$ ; $+ = 0.05 < p \leq 0.1$ ; $* = 0.01 < p \leq 0.05$ ; $** = 0.001 < p \leq 0.01$ ; and $*** = p \leq 0.001$ .

**Table 3.** Mean values and parametric/non-parametric (Mann-Kendall) temporal trends in monthly total precipitation, rainfall, snowfall, and end of month total snow on ground at the Kamloops A climate station between 1990 and 2011. Mean values are in units of mm except snow on ground (cm). Corresponding temporal trends are on a mm(cm)/yr basis.

| Descriptor | Jan | Feb | Mar | Apr | May | Jun | Jul | Aug | Sep | Oct | Nov | Dec |
| --- | --- | --- | --- | --- | --- | --- | --- | --- | --- | --- | --- | --- |
| Total precipitation |  |  |  |  |  |  |  |  |  |  |  |  |
| Mean | 25.8 | 13.2 | 17.2 | 14.1 | 33.0 | 42.4 | 33.4 | 22.9 | 26.6 | 24.8 | 29.2 | 29.5 |
| Trend |  |  |  |  |  |  |  |  |  |  |  |  |
| Parametric | NS | NS | 0.47/+ | NS | NS | -1.3/* | -1.3/+ | -1.4/* | NS | NS | NS | -1.0/+ |
| Mann-Kendall | NS | NS | NS | NS | NS | -1.6/** | NS | -1.1/+ | NS | NS | NS | NS |
| Monthly total of daily rainfall |  |  |  |  |  |  |  |  |  |  |  |  |
| Mean | 6.6 | 7.0 | 11.9 | 13.8 | 33.0 | 42.4 | 33.4 | 22.9 | 26.6 | 24.5 | 17.9 | 8.1 |
| Trend |  |  |  |  |  |  |  |  |  |  |  |  |
| Parametric | NS | NS | NS | NS | NS | -1.3/* | -1.3/+ | -1.4/* | NS | NS | NS | NS |
| Mann-Kendall | NS | NS | NS | NS | NS | -1.6/** | NS | -1.1/+ | NS | NS | NS | -0.42/+ |
| Monthly total of daily snowfall |  |  |  |  |  |  |  |  |  |  |  |  |
| Mean | 19.2 | 6.3 | 5.2 | 0.3 | 0.0 | 0.0 | 0.0 | 0.0 | 0.0 | 0.3 | 11.2 | 21.4 |
| Trend |  |  |  |  |  |  |  |  |  |  |  |  |
| Parametric | NS | NS | NS | NS | NS | NS | NS | NS | NS | NS | NS | NS |
| Mann-Kendall | NS | NS | NS | NS | NS | NS | NS | NS | NS | NS | NS | NS |
| End of month total snow on ground |  |  |  |  |  |  |  |  |  |  |  |  |
| Mean | 5.0 | 0.7 | 0.0 | 0.0 | 0.0 | 0.0 | 0.0 | 0.0 | 0.0 | 0.0 | 2.6 | 7.1 |
| Trend |  |  |  |  |  |  |  |  |  |  |  |  |
| Parametric | NS | NS | NS | NS | NS | NS | NS | NS | NS | NS | NS | NS |
| Mann-Kendall | NS | NS | NS | NS | NS | NS | NS | NS | NS | NS | n/a<sup>a</sup> | NS |

<sup>a</sup> Data missing for 2008.
NS= $p > 0.1$ ; += $0.05 < p \leq 0.1$ ; \*= $0.01 < p \leq 0.05$ ; \*\*= $0.001 < p \leq 0.01$ ; and \*\*\*= $p \leq 0.001$ .

**Table 4.** Mean values and parametric/non-parametric (Mann-Kendall) temporal trends in monthly mean hourly wind speeds, direction of the maximum monthly wind gust, and the speed of the maximum monthly wind gust at the Kamloops A climate station between 1990 and 2011. Mean values are in units of km/h except direction of the maximum monthly wind gust (°). Corresponding temporal trends are on a km/h(°)/yr basis.

| Descriptor | Jan | Feb | Mar | Apr | May | Jun | Jul | Aug | Sep | Oct | Nov | Dec |
| --- | --- | --- | --- | --- | --- | --- | --- | --- | --- | --- | --- | --- |
| Monthly mean of the hourly wind speeds |  |  |  |  |  |  |  |  |  |  |  |  |
| Mean | 12.0 | 11.9 | 13.3 | 12.8 | 11.8 | 11.1 | 10.6 | 10.0 | 10.0 | 11.6 | 13.2 | 13.0 |
| Trend |  |  |  |  |  |  |  |  |  |  |  |  |
| Parametric | NS | NS | 0.13/* | NS | NS | NS | -0.04/+ | NS | 0.10/* | NS | NS | NS |
| Mann-Kendall | NS | NS | NS | NS | NS | NS | -0.03/+ | NS | 0.12/* | NS | NS | NS |
| Direction of maximum monthly wind gust |  |  |  |  |  |  |  |  |  |  |  |  |
| Mean | 21.8 | 20.6 | 23.0 | 23.7 | 21.2 | 23.3 | 23.0 | 23.1 | 24.9 | 22.7 | 18.6 | 22.1 |
| Trend |  |  |  |  |  |  |  |  |  |  |  |  |
| Parametric | NS | NS | NS | NS | NS | NS | NS | NS | -0.37/+ | NS | NS | NS |
| Mann-Kendall | 0.33/* | NS | NS | NS | NS | NS | NS | NS | -0.15/* | NS | n/a<sup>a</sup> | NS |
| Speed of maximum monthly wind gust |  |  |  |  |  |  |  |  |  |  |  |  |
| Mean | 63.1 | 62.3 | 67.6 | 67.6 | 67.0 | 63.5 | 66.0 | 67.4 | 59.7 | 63.7 | 70.0 | 66.9 |
| Trend |  |  |  |  |  |  |  |  |  |  |  |  |
| Parametric | 0.9/* | NS | NS | NS | NS | NS | NS | NS | 0.8/* | NS | NS | NS |
| Mann-Kendall | 0.9/+ | NS | NS | NS | NS | NS | NS | NS | 0.9/* | NS | n/a | -0.6/+ |

<sup>a</sup> Data missing for 2008.
NS= $p > 0.1$ ; += $0.05 < p \leq 0.1$ ; \*= $0.01 < p \leq 0.05$ ; \*\*= $0.001 < p \leq 0.01$ ; and \*\*\*= $p \leq 0.001$ .

Mean hourly wind speeds and maximum monthly wind gust speeds are increasing (by ~ 1 km/h/decade) during September, with a corresponding shift in the wind direction of the maximum monthly wind gust (at a rate of 1.5 to 3.7°/decade) to a more northerly origin. However, it should be noted that being in an east-west trending valley, the predominant wind direction at Kamloops is from the east on an annual basis, from the west during May, June, and July, and from the east for the remainder of the year (Environment Canada 2012). The directions of the maximum hourly wind speed and the maximum gust are from the south (including southwest and southeast) between May and November and from the west (including northwest and southwest) between December and April.

A strong seasonal intervariable correlation inversion is clear between the 6 am average RH and the corresponding mean monthly temperature (MT; Table 5). Significant negative correlations exist between the 6 am average RH and MT between May though September, changing to significant positive correlations between these two variables from December to March, and no significant correlations during the transition periods of April and October/November. Significant negative correlations exist between the 3 pm average RH and MT between May though September and no significant correlations during the remainder of the year. Mean monthly wind speed (MWS) is negatively correlated with average RH from October to February (6 am) and November to January (3 pm). Significant positive correlations are evident between RH and MWS during March (3 pm) and July (6 am and 3 pm).

**Table 5.**
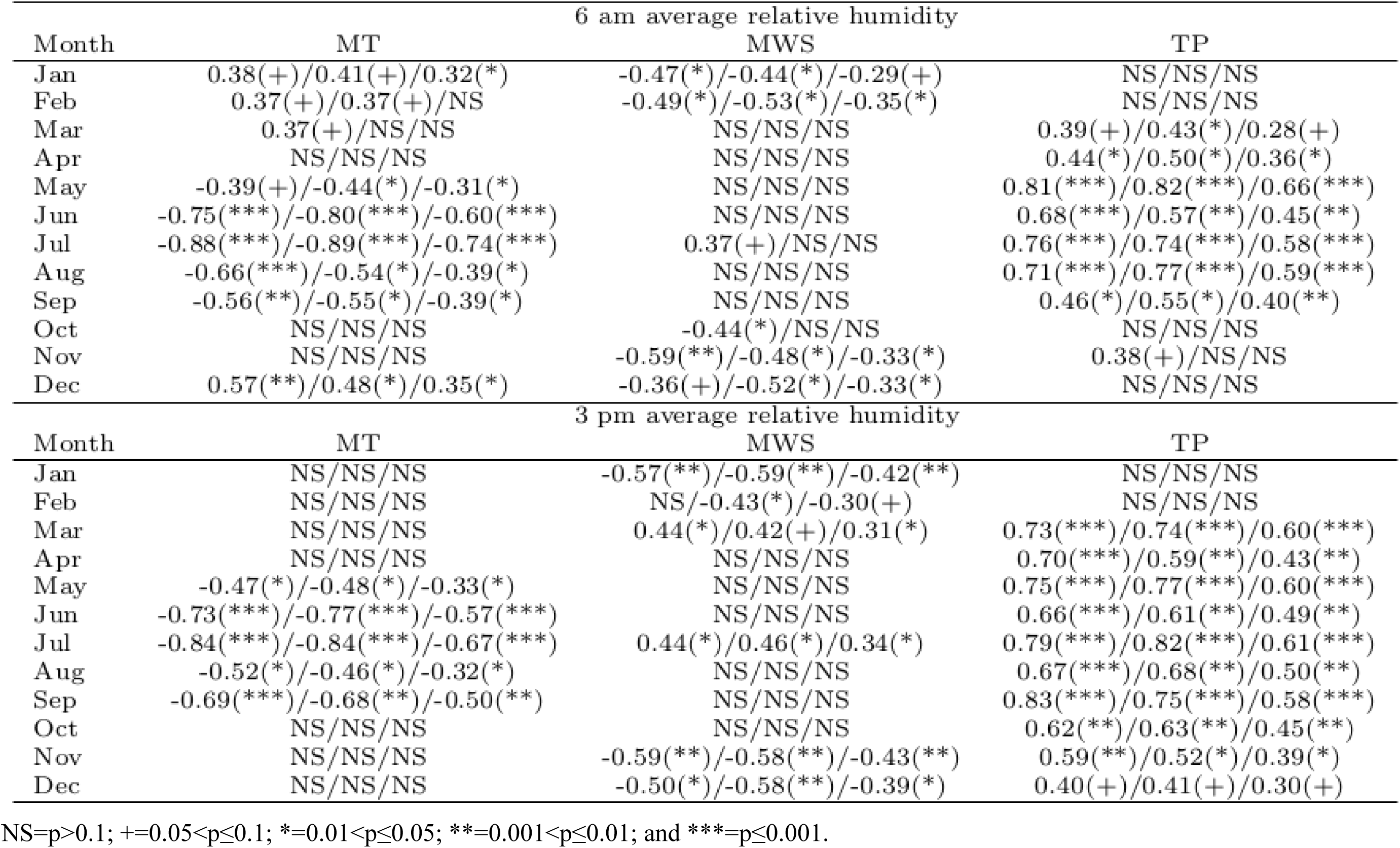
Correlation coefficients and level of statistical significance between the 6 am and 3 pm monthly average relative humidity and the corresponding mean temperature (MT), mean wind speed (MWS), and total precipitation (TP) at the Kamloops A climate station between 1990 and 2011. Values are presented as parametric correlation/Spearman non-parametric rank correlation/Kendall non-parametric rank correlation.

There are positive correlations between monthly total precipitation (TP) and the 6 am (March to September and November) and 3 pm (March to December) average RH for the majority of the year. Thus, warmer temperatures and reduced precipitation during the snow-free season lead to lower RH via the combined effects of greater evaporation in an arid region and the influence of warmer, dryer summertime high pressure weather patterns. Warmer temperatures during the winter period generally result in higher RH, likely from the combined effects of snowmelt induced humidification and the influence of warmer, humid winter-time low pressure weather patterns.

Recent work has shown a negative logarithmic relationship between summertime RH and corresponding dust generation at mine sites in central and northern Canada (Chang et al. 2012). For each 10% decline in RH (the approximate rate of RH decline observed at Kamloops during the summer months over the past two decades) between 40 and 90% RH, these authors reported an increase in mine dust concentrations behind haul trucks ranging from 1.3 and 2.4-fold. While site/activity specific differences in dust generation will exist between mine sites owing to variations in the particle size distribution, specific gravity, and hygroscopic nature of the various materials, the results indicate that if declining RH trends continue at Kamloops, current and proposed mine sites within the region are likely to generate progressively more dust (assuming no mitigative measures are undertaken) over time relative to the current climate. It is also important to note that lower RH (especially if coupled to higher temperatures) poses an increased risk of wildfires, a natural hazard of regular occurrence in the study area.

Overall, our findings indicate that the continuation of declining trends in RH near Kamloops, British Columbia, Canada, may increase the quantity of dust and other atmospheric particulate generation from both natural and anthropogenic sources, possibly resulting in additional threats to local and regional air quality, thereby necessitating inclusion in air quality management planning and modeling efforts.

